# Regulation between LRRK2 and PP2A signaling in cellular models of Parkinson’s disease

**DOI:** 10.1101/2025.10.07.680857

**Authors:** Panagiotis S. Athanasopoulos, Anna Memou, Franz Y. Ho, Ahmed Soliman, Henderikus Pots, Vassiliki Papadopoulou, Felix von Zweydorf, Saiganesh Sriraman, Alexander Marc Thouin, Laurine Vandewynckel, William Sibran, Marie-Christine Chartier-Harlin, R. Jeremy Nichols, Elisa Greggio, Jean-Marc Taymans, Christian Johannes Gloeckner, Hardy J. Rideout, Arjan Kortholt

## Abstract

Mutations in Leucine-rich repeat kinase 2 (LRRK2) are the most frequent cause of late-onset familial and idiopathic Parkinson’s disease (PD), known to date. Importantly, recent data from post-mortem tissue as well as biomarker studies suggest that independent of mutations, increased kinase activity of LRRK2 plays an essential role in idiopathic PD pathogenesis. Despite extensive research on LRRK2, its activation mechanism(s) and how the various mutations result in increased kinase activity and neuronal death is still not completely understood. Accumulating evidence points to LRRK2 phospho-regulation, both auto-phosphorylation and phosphorylation by other kinases, as one potential molecular trigger of its activation. LRRK2 activation and localization is regulated by phosphatases such as Protein phosphatase 1 (PP1) and Protein phosphatase 2A (PP2A), however the exact mechanism of this phospho-regulation is not known. Our data reveal that in vitro PP2A dephosphorylates sites within the RocCOR-GTPase domain of LRRK2 and as a result de-stabilizes LRRK2 dimers, with consequent reduction of its kinase activity. Strikingly, our data further highlight that LRRK2 in turn phosphorylates the catalytic subunit of the PP2A holoenzyme PPP2CA at its critical residue T304. Furthermore, LRRK2-mediated phosphorylation of PP2CA T304 alters the methylation of the C-terminus, which is crucial for both holoenzyme formation and catalytic activity. Importantly, expression of WT-PPP2CA protects from LRRK2-G2019S induced neuronal cell death, while PPP2CA-T304 mutants fail to do so, suggesting that impaired PP2A holoenzyme formation might be detrimental for LRRK2-PD.

## Introduction

PD is the second most common neurodegenerative disease after Alzheimer’s disease, affecting 1–2% of the Western world’s population older than 60 years [1]. From a pathological point of view, it is characterized by the progressive death of dopaminergic neurons in the ventral midbrain associated with the formation of fibrillar aggregates that are α-synuclein (αSyn) enriched in most forms of PD. Mutations in leucine-rich-repeat kinase 2 (LRRK2) have been identified to be the main cause of late-onset hereditary Parkinson’s disease (PD) [2]. From a clinical perspective, it is largely indistinguishable from the more common idiopathic PD; however, there are some neuropathological differences. LRRK2 is a large multidomain protein consisting of two enzymatic domains, namely a RocCOR GTPase domain, belonging to the Roco protein family of G proteins, which is followed by a Serine/Threonine kinase domain [3–7]. Several pathogenic point mutations located in the GTPase domain (such as N1437H, R1441C/G/H/S) and in the kinase domain (G2019S, I2020T), as well as the COR domain (Y1699C), all lead to increased kinase activity and subsequently play an important role in PD causality [8–10]. Moreover, LRRK2 kinase activity is not only increased in LRRK2 familial PD, but also in idiopathic patients in the absence of a mutation [11,12] suggesting that LRRK2 (dys)function plays a broader role in PD. Therefore, the regulation of LRRK2 kinase activity is considered a key therapeutic target in familial and sporadic PD.

Despite a vast amount of research and the identification of several LRRK2 substrates and interaction partners, the exact cellular function of LRRK2, the underlying mechanism that results in increased LRRK2 activity and how this subsequently results in the progression of the disease is still not completely understood [13]. Importantly, Rab proteins have been identified as both upstream regulators and as key physiological substrates of LRRK2 [14–16]. Accumulating evidence suggests that this LRRK2/Rab pathway functions at the interface of vesicular trafficking and autophagy. LRRK2 is able to phosphorylate a subset of Rab GTPases within their Switch-II motif thereby controlling interaction with effectors and cellular trafficking. The PPM1H phosphatase is able to dephosphorylate this motif in the substrate Rabs [17,18] thereby counteracting LRRK2 signaling.

Phospho-regulation plays an important role in the LRRK2 activation cycle [19–22]. The N-terminal LRRK2 phosphorylation cluster (S910, S935, S955, S973) regulates binding to 14-3-3, which is important for LRRK2 localization, activity and neurotoxicity [22–29]. Furthermore, phosphomutant studies at the heterologous phosphosites of LRRK2 (around S910 and S935) showed that phosphodead LRRK2 displays hyperactivatable kinase activity while phosphomimicking LRRK2 shows reduced kinase activity markers, pointing to a role in LRRK2 heterophosphosites in regulating LRRK2 kinase activation [22], however so far, the role of LRRK2 autophosphorylation sites in LRRK2 kinase activation and toxicity is unknown. Phosphorylation of LRRK2 is regulated by both upstream kinases and its own kinase domain. PKA has been reported to be an upstream regulator of S935 phosphorylation. [25,30,31]. Furthermore, the PD R1441C/G/H mutation also impairs PKA-induced S1444 LRRK2 phosphorylation, and disrupts its interaction with 14-3-3 [32]. Dephosphorylation of S910/S935/S955 and S973 is regulated by Protein Phosphatase 1 (PP1) and thereby stimulates the dissociation of 14-3-3 proteins from LRRK2 [26]. In addition, several auto-phosphorylation sites within the Roc domain have an impact on GTPase and kinase activity [7,32–38]. We have recently revealed an intramolecular feedback regulation of the LRRK2 Roc G domain by a LRRK2 kinase dependent mechanism [37,38]. Biochemical analysis showed that auto-phosphorylation of the Roc domain reduced the GTPase activity, most likely by regulating the monomer-dimer equilibrium of LRRK2 [37,38]. Furthermore, the phosphorylation status of the T1503 residue has been shown to be essential for the kinase activity as well as for the GTP binding properties of LRRK2 [36] while S1444 phosphorylation has been shown to influence the kinase activity of LRRK2 [32].

We have previously reported subunits of the PP1 and PP2A classes of phosphatases as well as the PAK6 kinase as regulators of LRRK2 dephosphorylation [26,39–43]. However, so far, no phosphatases have been identified that regulate the phosphorylation status of the Roc domain. We have previously identified PP2A as an interacting partner of the LRRK2 RocCOR-tandem domain [44]. Furthermore, activation and overexpression of PP2A results in neuroprotection in SH-SY5Y cells expressing LRRK2-R1441C, primary cortical neurons expressing LRRK2-G2019S, and in a LRRK2-*Drosophila* model [44,45]. In addition, we have shown that PAK6-mediated phosphorylation of the PP2A subunit PPP2R2C regulates recruitment of PPP2R2C to the LRRK2 complex and PPP2R2C subcellular localization [46]. Taken together, this shows that PP2A plays an important role in regulating LRRK2 activation and functioning.

In this study, we demonstrate that in vitro PP2A dephosphorylates LRRK2 at Threonine 1503 (T1503) of the Roc domain and thereby regulates LRRK2 dimerization and kinase activity. Most importantly, our data reveal that LRRK2 can in turn also phosphorylate Threonine 304 (T304) of the catalytic subunit of PP2A (PPP2CA). PP2A holoenzyme consists of three different subunits, the scaffold subunit, the catalytic subunit and the regulatory subunit. Depending on which regulatory subunit binds to the scaffold/catalytic core dimer complex, the holoenzyme is then localized to different compartments of the cell [47]. In addition, the methylation of the PPP2CA subunit is crucial for the holoenzyme complex formation. Interestingly, PPP2CA-T304 phosphorylation results in altered PPP2CA methylation, impaired PP2A holoenzyme assembly and reduced enzymatic phosphatase activity. Interestingly, PP2A activity is downregulated in alpha-synuclein PD models, having the same net-effect on PP2A activity as with hyper-active mutant LRRK2, and the methylation status of the C-terminal of PPP2CA plays an essential role in this regulation and progression of the disease. Understanding the regulation of PP2A, by post-translational modifications such as phosphorylation and methylation, can thus give important insight into the onset and progression of PD in general.

## Results

### PP2A dephosphorylates LRRK2 and regulates its kinase activity *in vitro* and in cells

To explain the previously observed neuroprotective properties of PP2A’s enzymatic activity in LRRK2-PD neuronal cell lines [44,45], it is crucial to characterize how PP2A impacts LRRK2 kinase function. It is clear that LRRK2 cycles between a monomeric cytosolic state that has low kinase activity and a high active dimeric membrane bound state[48]. In addition previous biochemical and structural work have confirmed that LRRK2 can exist as a monomer, dimer, and multimer[49]. The various available high resolution cryo-EM structures have shown that the RocCOR tandem domain is providing the main domain dimerization interface [50]. Consistently, constrained peptides targeting either the Roc or COR domain impair dimerization and inhibit LRRK2 kinase activity [51,52]. Since PP2A binds to the RocCOR domain, and the neurotoxicity of PD-mutant LRRK2 is tightly associated with its dimerization dependent kinase activity [48,51,53–57], we tested whether dimerization and kinase activity are altered by PPP2CA expression by using a method to label and purify dimeric LRRK2 from cells [57]. Over-expression of PPP2CA in HEK293T cells together with the proximity biotinylation LRRK2 construct pairs reduced LRRK2 dimerization (Fig. 1A). Moreover, the *in vitro* kinase activity of the isolated dimers (phosphorylation of NICtide) was also reduced in cells co-expressing WT-PPP2CA (Fig. 1B). The overexpression levels of WT-PPP2CA were also tested by western blotting (Supplementary Fig. 1). These data suggest that PP2A expression can regulate the dimerization and kinase activity of LRRK2. To further understand the mechanism by which PP2A regulates LRRK2 dimerization and kinase activity we next tried to identify the PP2A target sites. Previously, Liu et al., developed an assay in which they phosphorylated the human LRRK2 Roc domain using the kinase domain of the *Dictyostelium* LRRK2 orthologue Roco4 [58]. Since we have shown that the COR domain plays an important role in LRRK2 dimerization [51,52] we used a similar approach, but with purified recombinant purified LRRK2 RocCOR instead of a Roc-only protein [59]. The recombinant human RocCOR fragment of LRRK2 was incubated with recombinant purified Roco4 kinase in reaction buffer containing ATP. Consistent with Liu et al., we see that incubation of LRRK2 RocCOR with the Roco4 kinase domain results in phosphorylation of the RocCOR domain. Besides multiple Roco4 autophosphorylation sites, confirming that active kinase was used, we were able to detect significant phosphorylation at T1503 of the RocCOR fragment by a DIA-based MS approach (Supplementary Fig. 2).

**Figure 1:**
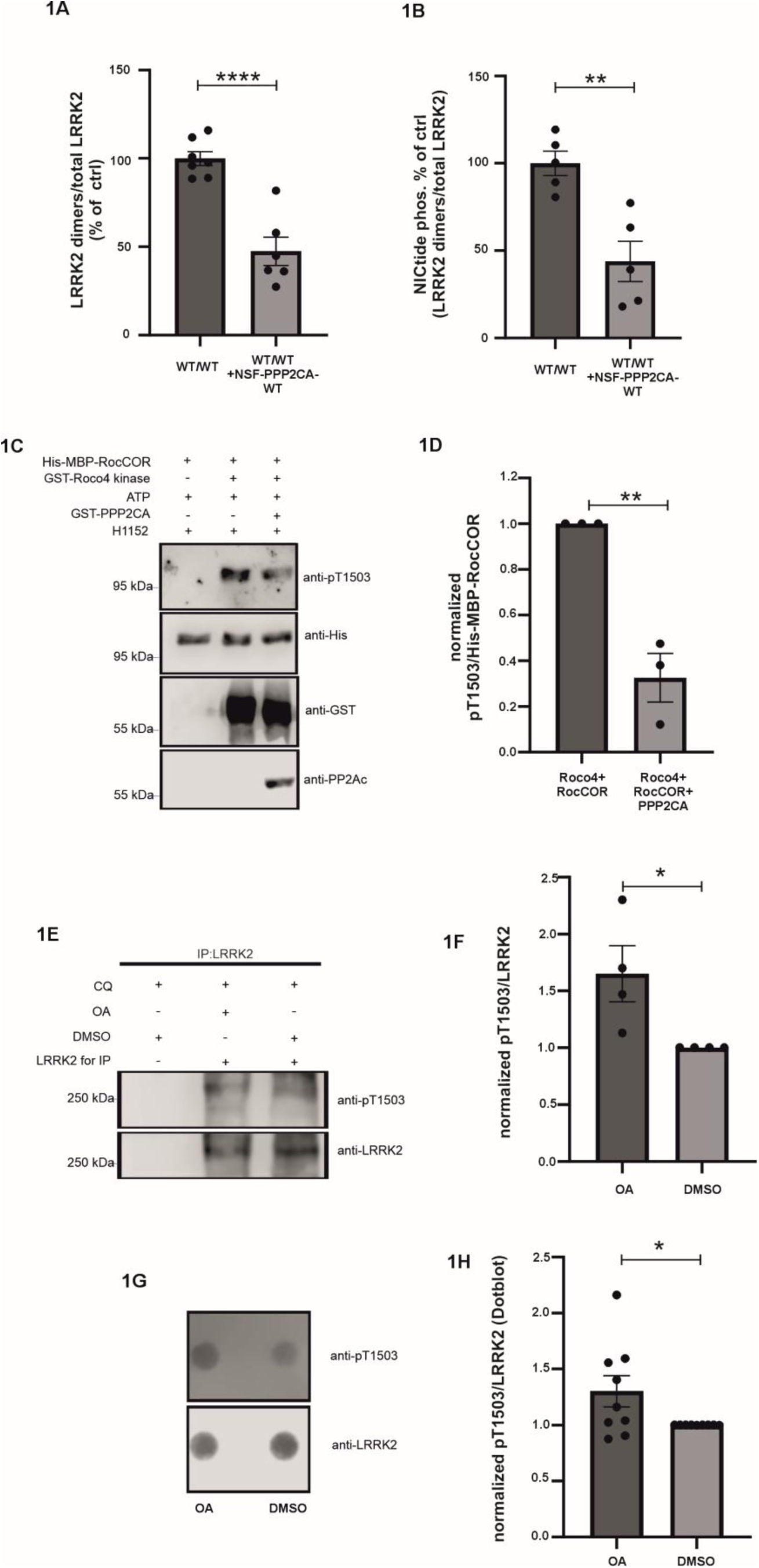
PPP2CA overexpression results in decreased LRRK2 dimerization and kinase activity, while PP2Ac is able to dephosphorylate LRRK2 *in vitro* and in cells at T1503. We examined the degree of dimer formation (A) and intrinsic kinase activity of isolated dimers (B) for wild-type (WT) LRRK2 co-expressed in HEK293T cells with WT-PPP2CA. Dimers were labeled with biotin *in situ* and captured on streptavidin-coated ELISA plates for ELISA and in-well activity measurements.. Regarding A), 7 biological replicates (n=7) and 6 biological replicates (n=6) were performed for WT/WT and WT/WT+NSF-PPP2CA-WT conditions respectively. For comparison, the mean of the WT/WT condition was set to 100%, and all other experimental conditions were expressed relative to this value. C) Purified His-MBP-RocCOR (aa 1293-1840) was incubated together with GST-Roco4 kinase and ATP to ensure RocCOR phosphorylation. The reaction was stopped by addition of the H1152 Roco4 kinase inhibitor and commercially available GST-PPP2CA was added. Phosphorylation of LRRK2 T1503 was determined by Western blotting (n=3 biological replicates). D) Quantification of (C). Shown is the normalized ratio of LRRK2 pT1503 phosphorylation (pT1503 signal divided by the His-MBP-RocCOR signal). E) LRRK2 pT1503 phosphorylation in A549 cells treated with DMSO or Okadaic acid for 60 minutes. In the presence of Okadaic acid the autophosphorylation of LRRK2 at T1503 is increased compared to cells treated only with DMSO (Okadaic acid solvent). As a control for the immunoprecipitation, no LRRK2 antibody was used for the immunoprecipitation (n=4 biological replicates). F) Quantification of (E). Shown is the ratio of the pT1503 and LRRK2 signal normalized to the DMSO condition. G) Dotblot from lysate of WT RAW264.7 macrophages treated with DMSO or Okadaic acid for 60 minutes prior to harvesting. H) Quantification of G). Shown is the ratio of the pT1503 and LRRK2 signal normalized to the DMSO condition (n=9 biological replicates). Data are means±SEM. *p < 0.05, **p < 0.01, ***p < 0.001, ****p < 0.0001, using unpaired T-test (two-tailed).

Since PP2A is interacting with LRRK2 via the Roc domain as a docking site [44], and T1503 residue has been shown to be important for the kinase and GTP binding properties of LRRK2, we tested with a commercial phospho-T1503 antibody, whether PP2A has an impact on LRRK2 phosphorylation status at this residue. Addition of the purified PPP2CA catalytic domain, results in significantly reduced phosphorylation of the RocCOR fragment of LRRK2 at T1503 (Fig. 1C + D). LRRK2 is highly expressed in lung and immune cells. A549 lung carcinoma cell line have high expression of LRRK2 and are therefore commonly used to measure the activation of LRRK2 in cells [60–65]. We used chloroquine-induced lysosomal stress to increase LRRK2 activity in A549 lung carcinoma cell line [63]. Additional treatment of A549 cells with Okadaic acid (OA), an inhibitor more selective for PP2A compared to PP1, resulted in the upregulation of the pT1503 signal from endogenously immunoprecipitated LRRK2, compared to the pT1503 signal of DMSO-treated (control) A549 cells (Fig. 1E + F). RAW264.7 macrophage cells were incubated with zymosan to stimulate LRRK2 activity and a dot-blot assay was performed to measure T1503 phosphorylation. Treatment of cells with zymosan has been shown previously to recruit LRRK2 to phagosomes and upregulate LRRK2 kinase activity [66,67], Consistent with our observations in A549 cells, the pT1503 signal in Okadaic acid RAW 264.7-treated cells was upregulated, compared to its DMSO-treated RAW 264.7 cells (control) (Fig 1G+ H). We next investigated whether the phospho-dead mutant version T1503A, also has an effect on LRRK2 activity and dimerization. Indeed, homo-dimers of T1503A-LRRK2 showed decreased dimerization and kinase activity, compared to wild-type LRRK2 (Fig. 2A + B); whereas hetero-dimers of WT/T1503A fail to de-stabilize dimers (Fig. 2C), yet have slightly reduced kinase activity (Fig. 2D). Although the T1503 residue is not directly located in the LRRK2 dimerization interface, it is located in close proximity to the nucleotide-binding site of LRRK2 [50] (Supplementary Fig. 3). Similarly other residues (T1343) which have been located to the RocCOR domain and in close proximity to the nucleotide-binding site and shown to affect the dimerization and the kinase activity of LRRK2 [37].Together our *in vitro* and cellular data suggest that PP2A can dephosphorylate LRRK2 at T1503 and thereby might regulate LRRK2 dimerization and kinase activity.

**Figure 2:**
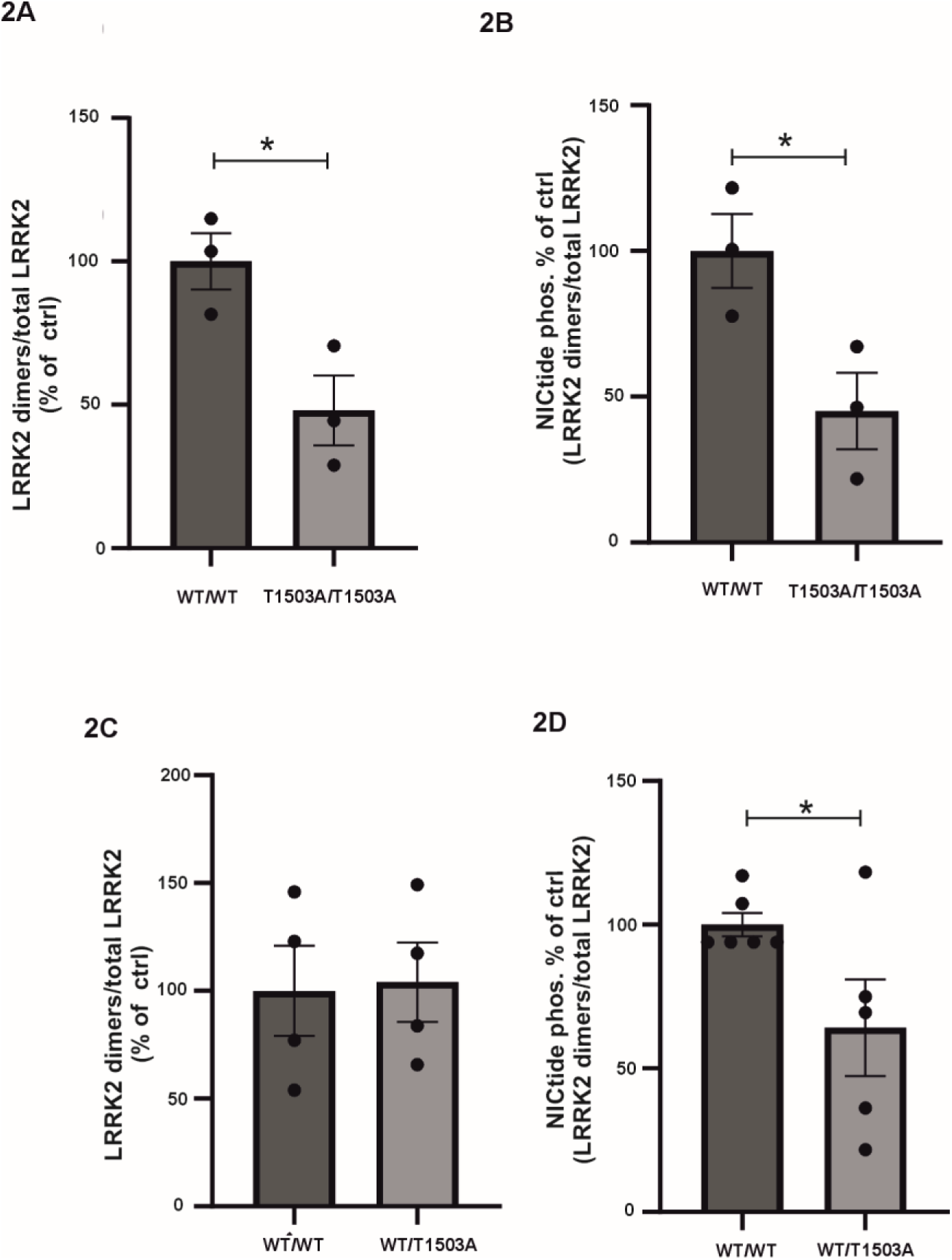
The phosphorylation status of the T1503 residue is essential for LRRK2 dimerization and kinase activity. To determine the impact of auto-phosphorylation at Thr1503-LRRK2 on dimerization and activity, we co-expressed proximity biotinylation construct pairs to isolate dimers of the following composition: WT/WT dimers, WT/T1503A hetero-dimers, or T1503A/T1503A homo-dimers. Dimer formation (2A+ C) and intrinsic kinase activity (2B+ D) were measured as above using an ELISA-based format (n=3 biological replicates for 2A and 2B, n=4 biological replicates for 2C. For 2D n=6 biological replicates were used for the WT/WT condition, and n=5 for the WT/T1503A condition). For comparison, the mean of the WT/WT condition was set to 100%, and all other experimental conditions are expressed relative to this value. Data are means±SEM. *p < 0.05, **p < 0.01, ***p < 0.001, ****p < 0.0001 using unpaired T-test (two-tailed).

### LRRK2 phosphorylates PPP2CA at T304 and thereby reduces its C-terminal methylation state

Surprisingly, using phospho-mass spectrometry we observed that LRRK2 can also phosphorylate PPP2CA at its critical T304 residue (phospho-Mass Spectrometry, Supplementary Fig 4). The T304 residue in the PPP2CA sequence is followed by a proline residue. Previous work showed that LRRK2 does have the potential to phosphorylate ThrPro sites [68]. Unfortunately, there is no commercial pT304-PPP2CA antibody available. Therefore, we used a pThrPro antibody to detect and confirm LRRK2-mediated phosphorylation of PPP2CA-T304. We co-expressed LRRK2-G2019S with either N-terminally twinstrep-FLAG (NSF) tagged WT-PPP2CA or the phosphodead T304A-PPP2CA mutant and measured the phosphorylation status after streptavidin pull down (Fig. 3A). We observed very clear phosphorylation of PPP2CA, while the T304A mutant showed a strongly reduced signal, confirming that we can use the pThrPro antibody to measure pT304. Next, purified human full length LRRK2 and PPP2CA were tested in an *in vitro* kinase assay (Fig. 3B). In the presence of ATP, LRRK2 is phosphorylating its auto-phosphorylation site S1292 and PPP2CA at T304, while in the presence of MLi-2, a potent and selective LRRK2 inhibitor, no phosphorylation was observed. To test LRRK2-mediated phosphorylation of PPP2CA under endogenous conditions, we immunoprecipitated the catalytic PPP2CA and PPP2CB subunits using a PP2Ac specific antibody. The pThrPro signal of the immunoprecipitated catalytic PP2A subunits was significantly reduced in the KO-LRRK2 RAW 264.7 cells, compared to WT RAW 264.7 (Fig. 3C + D). Consistently, the PP2A catalytic subunits are phosphorylated in A549 cells, however when the LRRK2 kinase activity was inhibited with MLi-2, the pT304 signal was diminished (Supplementary Fig. 5). Together this thus shows that PPP2CA is a LRRK2 substrate.

**Figure 3:**
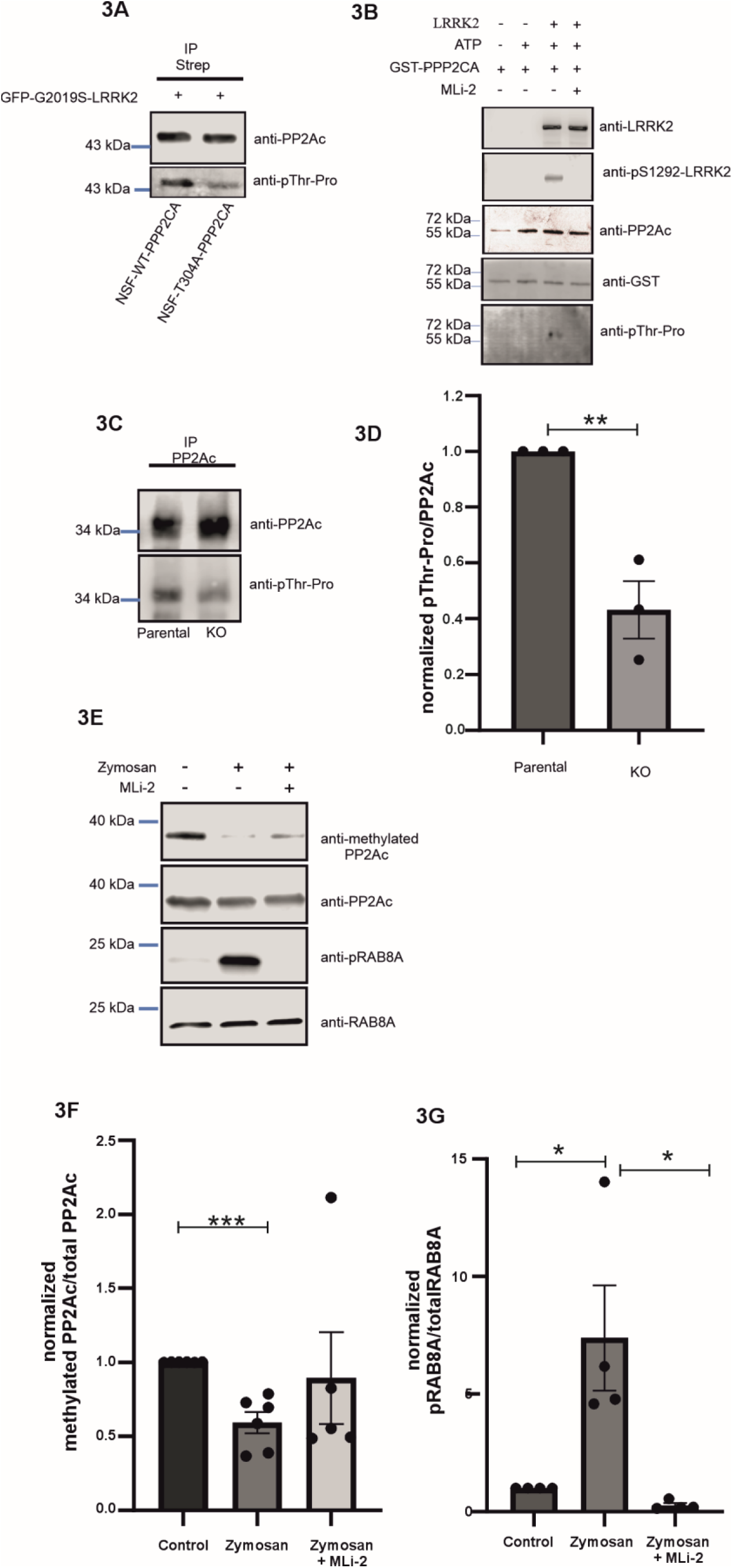
*In vitro* and *in cellulo* identification data showing LRRK2 phosphorylating PPP2CA at the Thr304 residue. A) GFP-LRRK2-G2019S was co-expressed with NSF-tagged WT-PPP2CA or T304A-PPP2CA in HEK293 cells. The PP2A proteins were isolated by streptavidin pull-down and phosphorylation of Thr304 WT-PPP2CA was visualized by a pThrPro specific antibody. B) Purified LRRK2 was incubated with commercially available purified WT-PPP2CA and ATP, with or without MLi-2 inhibitor (n=3 biological replicates). C) The catalytic PPP2CA and PPP2CB were immunoprecipitated from parental or KO-LRRK2 RAW 264.7 macrophages using the PP2Ac antibody. Phosphorylation was determined by western blotting (n=3 biological replicates). D) Quantification of panel C. Shown is the ratio of the pT304 and PP2Ac signal normalized to the WT condition. E) RAW 264.7 macrophages were stimulated with zymosan to achieve LRRK2 activation. RAB8A and phosphoRAB8A blots are depicted to demonstrate LRRK2 activation upon zymosan (in the presence or absence of MLi-2, inhibitor). F) Quantification of E) Shown is the ratio of methylated PP2Ac and total PP2Ac, normalized to the control (non-zymosan treated) condition (n=6 biological replicates for control and zymosan conditions, n=5 for zymosan+MLi-2 condition). G) Quantification of E) Shown is the ratio of phosphoRAB8A and total RAB8A, normalized to the control (non-zymosan treated condition) (n=4 biological replicates). Data are means±SEM. *p < 0.05, **p < 0.01, ***p < 0.001, ****p < 0.0001, using unpaired T-test (two-tailed).

Previous studies have shown that T304 phosphorylation alters the methylation of the C-terminus. Methylation of PPP2CA L309 removes the negative charge of the C-terminal carboxyl group, thereby allowing the interaction with the B regulatory subunits. Impaired methylation induces the dissociation of PPP2CA from the B regulatory subunits, resulting in changes in the subcellular localization of the PP2A holoenzyme and its substrate repertoire [69,70]. To address if activation of endogenous LRRK2 activation affects PP2A methylation we treated RAW264.7 macrophages with zymosan (Fig. 3E + F). Our data show that zymosan treatment results in LRRK2 activation as measured by RAB8A phosphorylation (Fig. 3E + G) and strongly reduced PP2Ac methylation (Fig. 3E + F). Consistently, vice versa inhibiting LRRK2 activity with MLi-2 results in increased, but not significant, increased levels of methylated PP2Ac) (Fig. 3E+ F). Together our data shows that LRRK2-mediated phosphorylation regulates the methylation status of PPP2CA.

### LRRK2 regulates PP2A holoenzyme formation and its catalytic activity

Previous data have shown that the C-terminal region of PPP2CA is essential for binding to its regulatory B subunit. C-terminally deleted PPP2CA mutants failed to interact with different regulatory B subunits ([69,70], Supplementary Fig. 6), while the phosphomimicking mutant T304D-PPP2CA, loses its interaction with PPP2R2B (B55β1) and PPP2R2A (B55α) subunits [70]. Consistently, our pull-down assays in HEK293 cell lysates showed that the interaction of PPP2R2B with T304D-PPP2CA is reduced compared to WT-PPP2CA or T304A-PPP2CA (Fig. 4A). Previous studies [44,45] have shown that PP2A is neuroprotective against mutant LRRK2-induced cell death; and that this is regulated via dephosphorylation of S6K and requires binding of PPP2R5C (B56γ3) PP2A to the holoenzyme [71]. Therefore, we next determined the interaction between PPP2CA phospho-mutants and PPP2R5C. Pull down experiments in HEK293 cells show that both T304A-PPP2CA and T304D-PPP2CA have strongly reduced interaction with PPP2R5C, compared to WT-PPP2CA (Fig. 4B + C). Therefore, we conclude that the de-regulation of T304 phosphorylation status of PPP2CA regulates holoenzyme formation. Next, we investigated whether T304A-PPP2CA or T304D-PPP2CA mutants exhibit alterations in their enzymatic phosphatase activity. By overexpressing and purifying WT/T304A/T304D-PPP2CA proteins from HEK293 cells and subsequently performing an *in vitro* phosphatase assay using a phosphopeptide and malachite green, we found that T304A-, as well as T304D-PPP2CA, exhibit lower phosphatase activity towards a phospho-Threonine peptide (K-R-pT-I-R-R), compared to the wild-type PPP2CA (Fig. 4D). Together, these data show that LRRK2 phosphorylates PPP2CA at position T304 and that this site is important for the regulation of holoenzyme complex formation and its catalytic activity.

**Figure 4:**
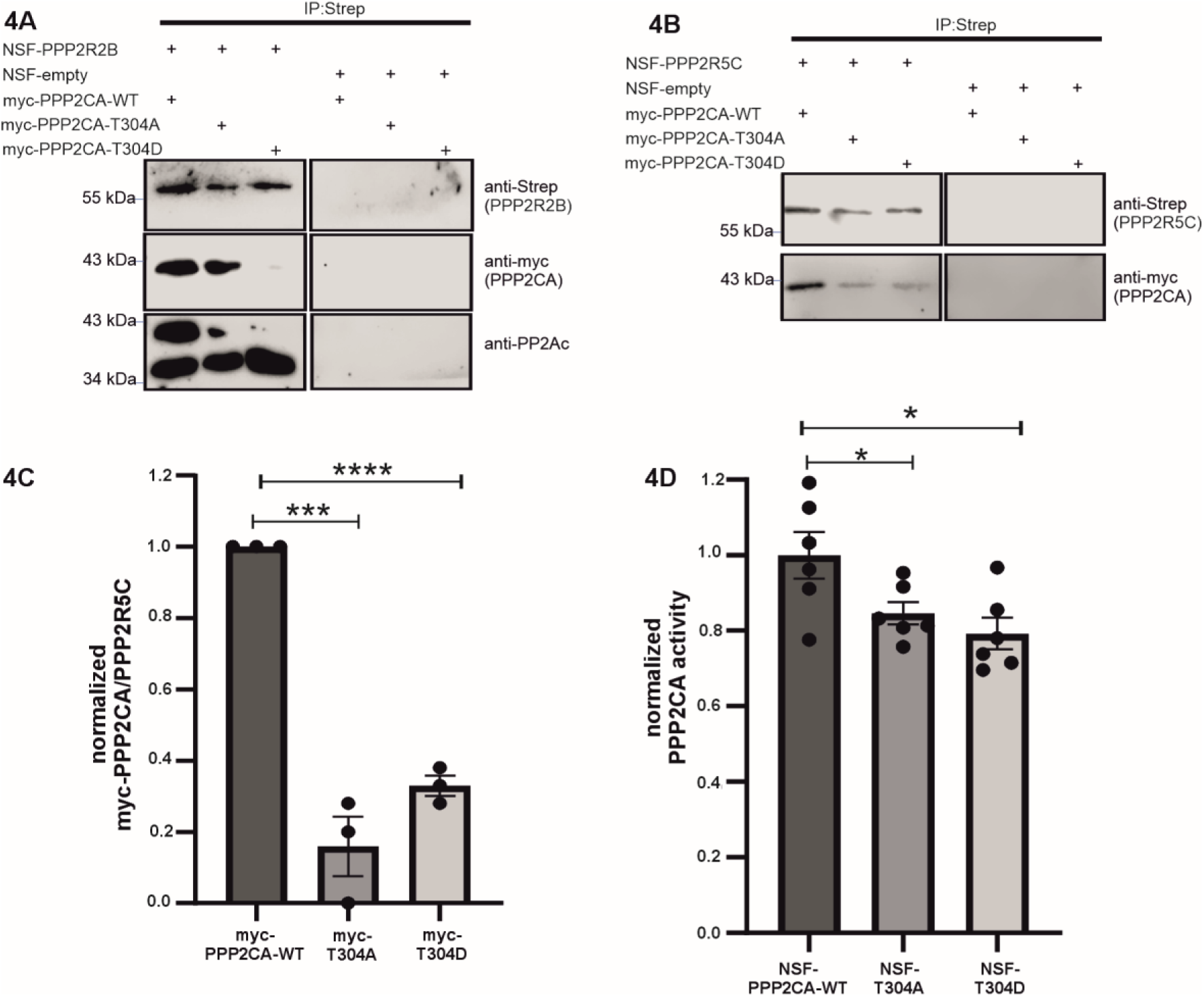
The regulation of the phosphorylation status of PPP2CA at T304 residue has an effect on the holoenzyme association as well as in its enzymatic activity. A) NSF tagged PPP2R2B or (B) NSF tagged PPP2R5C were co-expressed with the indicated myc-tagged PPP2CA constructs and the interaction was tested by immunoprecipitation. As a positive control, the endogenous catalytic PP2A subunit (anti PP2Ac staining, lower band, panel A), is interacting in the myc-PPP2CA-T304D sample pull down (n=3 biological replicates). C) Quantification of (B). D) PP2A activity of the indicated NSF tagged PPP2CA proteins was measured using an Ser/Thr Phosphatase assay kit (#17-127, Merck). The release of pmoles of phosphate is normalized to WT-PPP2CA. (n=2 for biological experiments with a triplicate per biological experiment). Data are means±SEM. *p < 0.05, **p < 0.01, ***p < 0.001, ****p < 0.0001 using unpaired T-test (two-tailed).

Previous studies have shown that either overexpression of PPP2CA or addition of sodium selenate, as a PP2A activator, was able to reverse G2019S-LRRK2-induced neurotoxicity in primary cortical neurons [44]. Similar data were shown in *Drosophila*, where overexpression of different single PP2A subunits were able to block G2019S-LRRK2-induced cell death in dopaminergic neurons [45]. To address the neurotoxicity of LRRK2, we first isolated primary mouse embryonic cortical neurons, and co-transfected them with LRRK2 (WT/G2019S) and PPP2CA-WT or the PPP2CA-T304A mutant. Immunofluorescence experiments confirmed co-expression of G2019S- LRRK2 and PPP2CA (WT or T304A) (Fig. 5A). Next LRRK2-induced cell death was measured, using a previously described assay that detects activated caspase-3 and fragmented nuclei, [51,52,72–74]. Consistent with previous studies, G2019S-LRRK2 overexpression resulted in enhanced active caspase-3/fragmented nuclei staining (Fig. 5B + C). As expected, the overexpression of WT-PPP2CA was able to protect from G2019S-LRRK2-induced neurotoxicity (Fig. 5C). In contrast, T304A-PPP2CA overexpression failed to do so (Fig. 5C). This suggests that phosphorylation of PPP2CA at this residue, and thus its phosphatase activity, is critical for its neuroprotective effects in this primary neuronal model.

**Figure 5:**
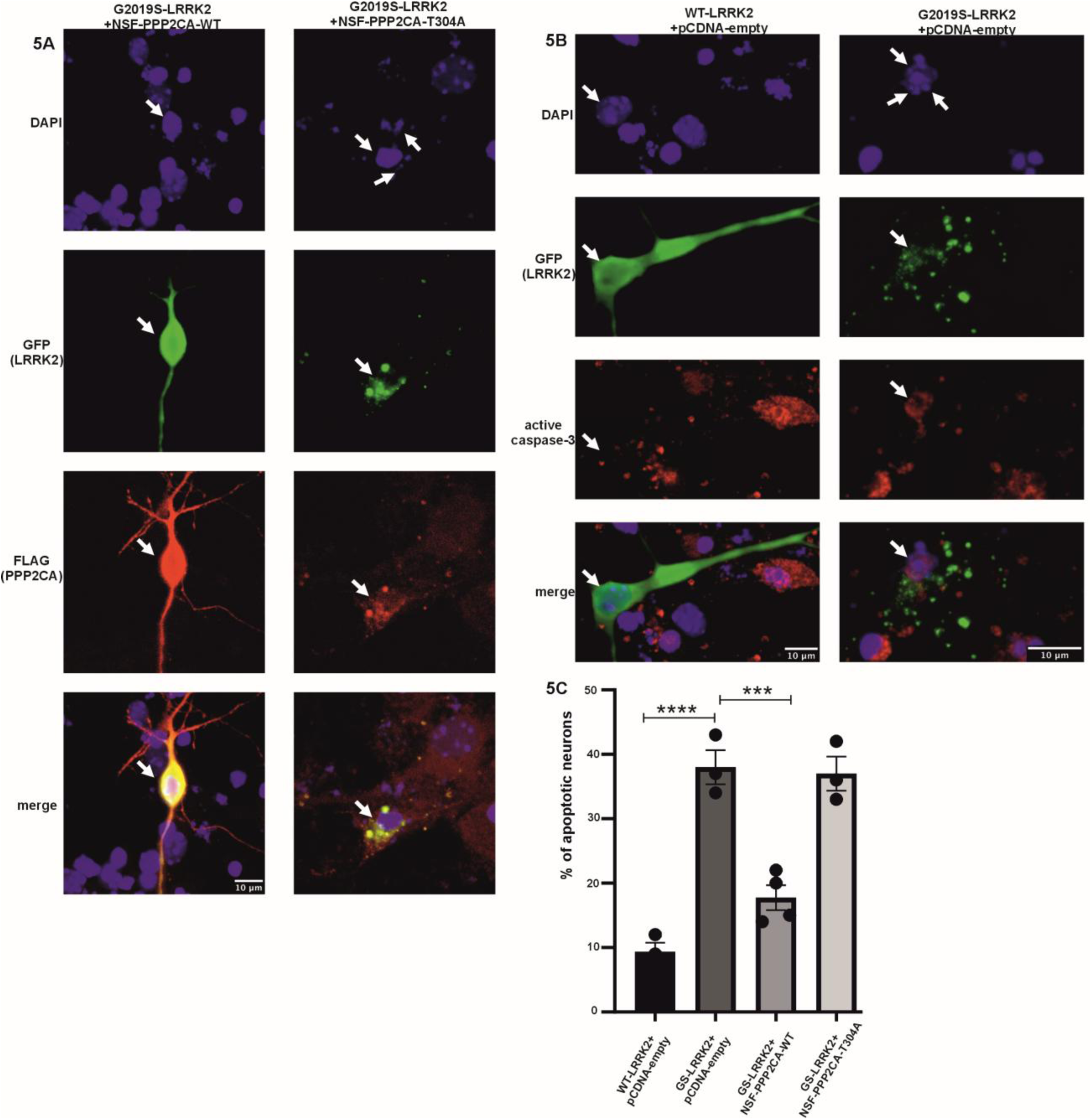
The overexpression of WT-PPP2CA, but not T304A-PPP2CA, rescues cortical neurons from G2019S-induced cell death. Primary embryonic mouse cortical neurons were transiently transfected with WT or G2019S-LRRK2 together with empty vector, WT-PPP2CA, orT304A-PPP2CA. Three days following transfection, the neurons were fixed and processed for immunofluorescence to label transfected cells. A) Representative image showing neurons being co-transfected with G2019S-LRRK2 (GFP reporter) and PPP2CA (Flag;red) WT or T304A, to confirm LRRK2/PPP2CA co-expression. DAPI staining has been used for nuclear morphology. B) Representative image of neurons being transfected with WT- or G2019S-LRRK2 (GFP reporter) co-stained with active caspase-3 (red) to test apoptosis levels. DAPI staining is also present in order to access fragmentation of nuclei (apoptosis) upon LRRK2 expression. C) The percentage of LRRK2/PPP2CA transfected neurons with apoptotic features was determined in a blinded manner. The plot shown is representative of 2-3 independent cultures and transfections, and each bar is comprised of at least 3 technical replicates (individual coverslips, with a minimum of 100 neurons counted in each coverslip) The images shown are image stacks recombined. The values represent the mean ±SEM, *** p<0.001; **** p<0.0001, F=36.84 with 4 DF, using one-way ANOVA, with Tukey’s post-hoc multiple comparisons test.

## Discussion

The importance of phosphatases in mutant LRRK2-induced parkinsonism has already been established by numerous studies. PPP1CA regulates the important N-terminal phosphorylation cluster, including S910 and S935. Interestingly, while PPP2CA alone does not have a significant direct effect on regulating S910/S935-LRRK2 phosphorylation [26], a phosphatome wide RNAi study revealed that PPP2CA associated to its regulatory subunit PPP2R2A/B was a potent regulator of those sites [43]. Related to this, LRRK2 autophosphorylation is reported to be sensitive to okadaic acid, suggesting that PP2A is also involved in regulating LRRK2 autophosphorylation sites [42]. In addition, PPM1H has been shown not to dephosphorylate LRRK2 directly, but to counteract LRRK2 function, by dephosphorylating LRRK2’s downstream targets, namely the small Rab GTPases [17]. Although we previously have shown that PP2A can bind to the ROC domain and that PP2A has neuroprotective effects in mutant LRRK2-induced cell death in SH-SY5Y neuroblastoma cells [44], and primary neurons (the present study), the molecular mechanism by which PP2A regulates LRRK2 signaling remained unclear.

Using *in vitro* and cellular models, we could identify the T1503 residue as one of the phospho-sites in the RocCOR domain of LRRK2, which can be dephosphorylated by PP2A. Both the LRRK2 T1503A mutant and overexpression of the catalytic subunit of the PP2A holoenzyme, PPP2CA, resulted in decreased LRRK2 dimerization and kinase activity. This suggests that the specific autophosphorylation-sites in LRRK2 modified by PP2A can regulate dimer stability, and by extension – kinase activity in vitro. Surprisingly, our data additionally revealed that LRRK2 could also phosphorylate PP2A in its catalytic subunit at the T304 residue. Furthermore, LRRK2-mediated PPP2CA T304 phosphorylation impairs holoenzyme formation and reduced phosphatase activity. Importantly, expression of WT-PPP2CA protects from G2019S-LRRK2 induced neuronal cell death, while the T304A-PPP2CA mutant fails to do so. Interestingly, PP2A activity is downregulated in αSyn PD models, having the same net-effect on PP2A activity as with hyper-active mutant LRRK2 [75,76]. Furthermore, in postmortem brain samples (substantia nigra-SN and frontal cortex (FC) from PD or dementia with Lewy body (DLB) patients, PP2A methylation in PD and DLB samples was greatly decreased, while Leucine carboxyl methyltransferase (LCMT-1), PP2A’s methylating enzyme, was significantly reduced in the SN of PD patients and protein phosphatase methylesterase (PME-1), a demethylating enzyme of PP2A, was elevated in the SN and FC of PD and DLB patients [75]. In addition, impaired activation of the PP2A phosphatase has been linked with autosomal recessive early-onset parkinsonism with intellectual disability [77].

Together, this shows that the regulation of the C-terminal segment of PPP2CA is crucial for the progression of PD in general. In addition to acting on LRRK2, PP2A has been shown to dephosphorylate αSyn at S129, which is believed to play a role in the pathogenesis of αSyn PD [78,79]. Moreover, it has been shown that PP2A can also dephosphorylate Tau *in vitro* [80–83]. Tau plays a crucial role in AD pathology and in addition has been identified as a potential PD factor in Genome wide associate studies [84]. Interestingly, it has been shown that PP2A activity is downregulated in AD brains [85,86]. Understanding the regulation of PP2A, by post-translational modifications such as phosphorylation and methylation, can thus give important insight into the onset and progression of PD in general.In summary, our data show that PP2A at least in vitro can dephosphorylate LRRK2, leading to de-stabilization of LRRK2 dimers and a reduction in its kinase activity. Importantly, LRRK2 can also phosphorylate the PP2A catalytic subunit. This latter event results in reduced PPP2CA methylation and impaired PP2A holoenzyme formation leading to the reduction of PP2A dephosphorylation activity. Our data thus also suggests that boosting PP2A activity by stimulating complex formation might be beneficial for mutant LRRK2-mediated PD and PD in general.

## Materials and Methods

### Cell Culture, pharmacological treatment, transfection and cell lysis

HEK293 cells were cultured in DMEM high glucose (#11960-044, Gibco), supplemented with 10% fetal bovine serum (FBS) (A31604-01, Gibco) and penicillin/streptomycin/glutamine (#10378-016, Gibco). HEK293 cells were transfected with jetPEI (#101-10, Westburg) according to the manufacturer’s protocol, or by incubating with DNA: CaPO4 precipitates. Protein expression was validated 48hrs after transfection with fluorescence microscopy and/or western blotting. Murine RAW 264.7 cells, parental (#sc-6003, ATCC) or Knock-out (KO) (# sc-6004, ATCC), were cultured in DMEM high glucose (#20-2002, ATCC), supplemented with 10% fetal bovine serum (FBS) (#20-2020, ATCC) and penicillin/streptomycin (#15070-063, Gibco). A549 cells were obtained from Prof. Amalia Dolga, (University of Groningen, the Netherlands) and were cultured in RPMI-160 (#12-702F, Lonza), supplemented with 10% FBS (#A31604-01, Gibco), and penicillin/streptomycin (#15070-063, Gibco). As a general lysis buffer protocol, cells were washed three times with wash buffer (50 mM Tris-HCl 7.5, 150 mM NaCl, 1mM EDTA, 2.5 mM EGTA). Then they were lysed (wash buffer supplemented with 0.1% NP-40, protease inhibitors (p2714-1btl, Sigma) [87], and phosphatase inhibitors (1mM Na3VO4, 1mM NaF, 5 mM β-glycerol phosphate).

In order to increase LRRK2 kinase activity and subsequently its autophosphorylation, 50 μM of chloroquine (#C6628, Sigma Aldrich) was used for treatment of A549 cells overnight [88]. For inhibition of the endogenous PP2A holoenzyme, 100 nM of okadaic acid (#O7885, Sigma Aldrich) was used according to the Lobbestael et al. 2013 study, [26] while DMSO was used as a vehicle control, 60 minutes before the cells got harvested.

### Dot-blotting

For dot-blotting Bio-Dot® (ser.nr. 84BR 33049, Biorad) device and nitrocellulose membranes 9x12 cm, 0.45 μm (#1620117, Bio-rad) were used. RAW264.7 macrophages were treated with 100 nM of okadaic acid, while DMSO was used as a vehicle control, 60 minutes before the cells got harvested. Approximately 0.5 μg per sample was loaded. Image J studio 6.0 has been used for the quantification of the signal intensity of the samples.

### Primary Antibodies

We used the following antibodies in this study: anti-PP2Ac (detects PPP2CA and PPP2CB proteins) (#2038, Cell Signaling, 1/1000 dilution working dilution), anti-pThreonine-Proline (#9391, Cell signaling, 1/250 working dilution), anti-pS1292-LRRK2 (#203181, Abcam, 1/500 working dilution), anti-GST (#27457701, GE Healthcare, 1/2000 working dilution), anti-His (#ab1187, Abcam, 1/100 working dilution), anti-LRRK2 (clone c41-2, Abcam, 1/1000 working dilution), anti-Strep (#34850, Qiagen, 1/5000 working dilution), anti-myc (#9E10:sc-40, Santa Cruz Biotechnology, 1/1000 working dilution), and anti-pT1503 (#154423, Abcam, 1/500 working dilution). For the detection of the C-terminal part (Leu309) of PP2Ac methylation, the following antibody (methyl-PP2A-Cα/β Antibody 2A10: #sc-81603, Santa Cruz Biotechnology, 1/500 working dilution) was used. Phospho RAB8A antibody (#ab230260, abcam, 1/1000 working dilution), and RAB8A antibody (#6975, Cell Signalling,1/1000 working dilution) were used for validation of LRRK2 activity.

### Purification of dimeric LRRK2

To purify LRRK2 dimers to be used in our *in vitro* assessments of kinase function, we relied on the proximity biotinylation technique previously described [57]. Briefly, two cDNAs were created encoding LRRK2 fusions with biotin ligase (BirA; N-term, Flag-tagged) and an acceptor peptide (AP, N-term; c-Myc tagged), and over-expressed in HEK293T cells grown in biotin-depleted medium (OptiMEM+2% FBS). After 48hr following transfection, the cells were extensively washed in PBS, given a brief biotin pulse (50 μM, 5 min, 37°C), followed by another 3X washes in PBS, centrifuged and the pellet snap frozen in a dry-ice/MeOH bath. Following lysis, extracts were diluted in TBST/BSA (10 mM Tris HCl, pH 7.6; 100 mM NaCl; 0.1% Triton X-100; 1% BSA) and 2.5 μg of protein loaded in parallel ELISA plates, coated with streptavidin (SA; to capture biotinylated LRRK2 dimers) and anti-LRRK2 (to quantify LRRK2 over-expression). To detect and quantify dimeric LRRK2, SA-coated plates were incubated with HRP-conjugated anti-Flag antibodies (1hr, room temperature). Since the biotin tag is only present on AP-LRRK2 fusions, and the Flag epitope tag is located on the BirA-LRRK2 fusion, by using HRP-Flag as our detector reagent, we are specifically labeling dimeric LRRK2 present in the ELISA plates. On parallel anti-LRRK2 coated plates (clone c41-2), total over-expressed LRRK2 is quantified using HRP-LRRK2 antibodies (clone N241), and used to normalize the relative amounts of dimeric LRRK2. We assessed the following LRRK2 dimers: WT/WT and G2019S/G2019S homo-dimers; as well as LRRK2 with the T1503 site mutated to Ala, in hetero- and homo-dimer conformations. We co-expressed PPP2CA (WT, T304A, or T304D) with WT or mutant G2019S-LRRK2 dimer construct pairs. Here, the ratio of expression of LRRK2 to PPP2CA was maintained at 3:1. Dimeric LRRK2 was determined as above with the following modification: LRRK2 dimers were estimated in SA-coated ELISA plates using HRP-conjugated anti-LRRK2 (clone N241; NeuroMab). We measured *in vitro* kinase activity of captured dimers using NICtide as a peptide substrate as described previously [57]. To assess phosphorylation at Ser935 in dimeric LRRK2, we used anti-pS935 LRRK2 conjugated to HRP (Expedion/Abcam) as our detector reagent, as recently described [89].

### Zymosan cell treatment and Lysis buffer for the detection of PP2Ac methylation

RAW 264.7 cells were stimulated with Zymosan (#Z4250, Sigma) for 30 min (20 particles per cell) prior to cell harvesting. RAW cells were also treated with 1µM of MLi-2 for 90 minutes prior to cell harvesting. In order to detect the methylation levels of the PP2Ac C-terminal, we followed the protocol as suggested by Yabe et al., [90]. In more detail, RAW264.7 macrophage cells were lysed in a buffer containing 50 mM Tris/HCl (pH 8.0), 5 mM EDTA, 5 mM EGTA, 1% Triton X-100, 1 mM Na3VO4, 20 mM sodium pyrophosphate, and Roche Complete protease inhibitor mixture. For measuring PP2Ac methylation level, 1 µM of ABL127 (#SML0294 Sigma-Aldrich) was added in a lysate buffer. To validate that zymosan increases LRRK2 activation, and to demonstrate elevation of phosphorylation of its physiological downstream RABs substrates, we added 0.1 µg/ml Microcystin-LR (#ALX-350-012, Enzo Life Sciences) and 0.5mM Diisopropylfluorophosphate/DIFP (#D0879, Sigma)[91].

### Preparation of primary mouse neuronal cultures, and assessment of neuronal death

Primary cells were prepared in the Animal Care Facilities of the Academy of Athens. Animal breeding and handling were performed in accordance with the European Communities Council Directive 86/609/EEC guidelines, and all animal procedures were approved by the Academy of Athens Institutional Animal Care and Use Committee. Embryonic day 16 (E16) pregnant C57BL mice were used in this study, with primary cortical neurons prepared as described [72,92]. The pregnant mice were sacrificed by rapid cervical dislocation. Under aseptic conditions, cortices were removed and cut into small pieces before enzymatic digestion (trypsin 0.05%) and mechanical dissociation. Cells were centrifuged and counted and plated at a density of 150,000/cm^2^ in BrainPhys neuronal culture medium (StemCell Technologies) supplemented with SM1 Neuronal Supplement (StemCell Technologies), L-glutamine (0.5 mM) and penicillin/streptavidin. After 3-4 DIV, neurons were transfected using Lipofectamine 2000 (ThermoScientific) as per the manufacturer’s instructions. Neurons were transfected with WT or mutant (G2019S) human LRRK2 (in pcms-EGFP vectors), with EGFP co-expressed under a separate promoter. Co-transfections with WT, T304A, or control plasmid, were performed at a ratio of 3:1 with excess LRRK2 plasmid. After three days following transfection, the coverslips were washed in PBS and fixed in 3.7% paraformaldehyde for 20 min at 4°C. The neurons were processed for immunofluorescence labeling with the following antibodies: GFP (chicken; Abcam, 1/1000 working dilution), Flag (M2 mouse; Sigma-Aldrich, 1/200 working dilution), active caspase-3 (rabbit; R&D Systems, 1/1000 working dilution), and DAPI nuclear stain (1 µg/ml). Mounted coverslips were imaged on a Leica TSP5 multi-photon confocal microscope, and the Z-stacks processed in ImageJ, and Adobe Photochop. For quantification of apoptotic neuronal profiles, we used the approach described by Antoniou and colleagues [72].

### Immunoprecipitation of endogenous PPP2CA/ LRRK2 and streptavidin pull down

To purify the endogenous PP2A catalytic subunits from RAW264.7 cells, the anti PP2Ac antibody (#2038, Cell signalling), detecting both PPP2CA and PPP2CB, p, together with Protein A-conjugated magnetic beads (#161-4013, Biorad). Expression of endogenous PP2Ac between parental and KO RAW264.7 cells was similar (https://zenodo.org/records/19234971). For the immunoprecipitation of endogenous LRRK2 from A549 cells, the Abcam LRRK2 antibody (#c41-2, Abcam) was used with Protein A-conjugated magnetic beads. In both cases, the following lysis/wash buffer was used: 10 mM Tris-HCl pH 7.5, 150 mM NaCl, 0.5 mM EDTA, 0.5 % NP-40. To purify overexpressed NSF-PPP2CA, WT and T304A, as well as NSF-PPP2R2B and NSF-PPP2R5C, the cells were lysed with 50 mM Tris-HCl 7.5, 150 mM NaCl, 1mM EDTA, 2.5 mM EGTA, 1% NP-40, protease inhibitors (p2714, Sigma) and phosphatase inhibitors (1mM Na3VO4, 1mM NaF, 5 mM beta glycerol phosphate) and incubated with streptavidin beads (#2-1206-025, IBA Lifesciences) for 4 hrs. Finally, they were washed 3 times with lysis buffer. Overexpression of the PP2Ac subunits (NSF/myc-PPP2CA-WT, -T304A, and -T304D) was confirmed by western blotting (expression levels for the different constructs was similar) (https://zenodo.org/records/19234971).

### Phosphatase activity assay

Phosphatase activity of purified NSF-PPP2CA (WT and T304A) was measured using the Ser/Thr Phosphatase assay kit (#17-127, Merck; Fig. 4D). Over-expressed PPP2CA proteins were purified on streptavidin beads as described above, and the activity determined with proteins bound to the beads. The protocol for the phosphatase assay and representation of the data has been previously described [93].

### Purification of proteins, *in vitro* assays and western blot assay

The lysis buffer used for LRRK2 purification from HEK293 cells consisted of 30 mM Tris pH 8.0, 150 mM NaCl, 5 mM MgCl2, 3 μM DTT, 5 % glycerol, 0.1 mM nucleotide, 0.5% NP-40), while the wash buffer is similar to the lysis buffer (excluding NP40). Finally, the streptavidin beads were used (#2-1206-025, IBA Lifesciences), while the 10x elution buffer (#2-1000-025, IBA Lifesciences) was diluted in wash buffer. The RocCOR domain of human LRRK2 fused with His and MBP tags at the N-terminus was expressed in *E. coli* BL21 (DE3). Cells were lysed in buffer consisted of 50 mM HEPES (pH 8), 150 mM NaCl, 10 mM MgCl2, 10% Glycerol, 0.5 mM GDP, 5 mM β-mercaptoethanol and complete EDTA free protease inhibitor cocktail by sonication, and then purified using MBP-Trap affinity chromatography column (GE Healthcare). The protein was eluted in the presence of 10mM maltose. For the *in vitro* kinase-phosphatase assay, the reaction buffer consisted of 25 mM Tris pH 7.5, 30 mM MgCl2, 150 mM NaCl, and 2mM DTT. 18 µg *Dictyostelium discoideum* 18 µg GST-Roco4-kinase (purified as previously described in [94]) 4 µg of His-MBP-RocCOR (aa 1293-1840) and 50 µM ATP were incubated at 25 degrees for 30 minutes. In order to block the Roco4 kinase activity and subsequently the phosphorylation of the RocCOR fragment, 10 mM of H1152 (#2414, Tocris Bioscience) was added. Finally, 1 µg of commercially available GST-PP2Ac (#128557, Abcam) was added to the solution for 105 minutes. The buffer of the *in vitro* kinase assay (Fig. 3B) consisted of 25 mM Tris pH 7.5, 15 mM MgCl2, 150 mM NaCl, 2 mM DTT. Recombinant purified full-length wild-type LRRK2 (3 µg) [95], 0.5 µg commercially available GST-PPP2CA (#128557, Abcam), and 50 µM ATP were incubated at 25°C for 30 minutes. In one of the samples, 1µM MLi-2 (#5756, Tocris) was included as a positive control for LRRK2 kinase inhibition. For western blotting, the wet transfer protocol has been used while nitrocellulose membrane 0.45 μm (#10600002, GE Healthcare Life science) or PVDF membrane 0.45 μm (#IPVH00010, Merck) was used. For band intensity measurement, ImageJ and Image studio 6.0 were used.

### Mass spectrometry

Phosphopeptide enrichment followed by LC-MSMS has been performed as previously described [96]. Briefly, after *in vitro* phosphorylation, protein samples were precipitated with chloroform methanol and re-dissolved in 100 mM ammonium bicarbonate. The proteins were subsequently alkylated by DTT/ Idoacetaminde treatment and submitted to tryptic proteolysis over-night. The phosphopeptides have been enriched by TiO2 prior to their analysis on an Orbitrap Fusion mass spectrometer (Thermo-Fisher Scientific) coupled to a nanoflow HPLC system (Ultimate 3000 RSLC; Thermo-Fisher Scientific) which was equipped with a µPac C18 reverse phase column (Thermo-Fisher Scientific). Full scans were acquired at 120K resolution (mass range: m/z 400-1000). MS2 Spectra were acquired via DIA in the Orbitrap (15K resolution), using window-wise CID fragmentation in 31 m/z windows of 20 Da width (1Da overlap) and 2h gradients. For spectral library creation, LRRK2-G2019S was pre-phosphorylated *in vitro* and processed as described above and analyzed via DDA to generate a spectral library. Instead, Roco4 MS2 spectra were directly extracted from the DIA files by DIAUmpire (2.0) [97]. For this purpose, MS2 spectra were searched in Mascot (v.2.5.1)[97] using carbamidomethyl cysteine as fixed modification and oxidizes methionine and phospho-STY as variable modifications. The search was performed against the human subset of the SwissProt database (release 2016_08) supplemented by the Roco4 sequence (total: 20178 entries). The peptide tolerance was set to 10 ppm, the MSMS tolerance was set to 20 mmu. Library creation from Mascot .dat files and final data analysis was performed in Skyline (v21.2.0.568) [98].

## Data availability statement

Raw data of the unedited western blot images are available in: https://zenodo.org/records/19234971.

## Author contributions

P.S.A performed most of the biochemical and cellular experiments with the help of S.S. and A.M.T.; A.M. H.J.R. and V.P. performed the dimerization assays, kinase assays and primary neuronal cell death assays; A.S. and H.P. assisted P.S.A. with cloning of different constructs; F.Y.H. assisted with the purification of proteins; F.v.Z. and C.J.G. performed the Mass spectrometry experiments/analysis; L.V., W.S., M.C.C.H, R.J.N., E.G. and J.M.T. interpreted experiments; P.S.A., A.K. and H.J.R. conceived the study; A.K. and H.J.R. supervised the experiments; P.S.A., A.K., and H.J.R. wrote the manuscript with the help and input of all other authors.

## Acknowledgments

We gratefully acknowledge Prof. Chi-Wu Chiang (National Cheng Kung University, Taiwan) for providing us with the PPP2CA, PPP2R5C and PPP2R2B sequences. We thank the staff of the Core Facility for Medical Proteomics at the University of Tubingen for technical assistance. We thank Nora Migdad for the structural analysis.

## Declaration of generative AI use

The authors declare that they haven’t used generative AI and AI-assisted technologies (AI tools) for the preparation of the manuscript.

---

**Supplementary Fig. 1:**
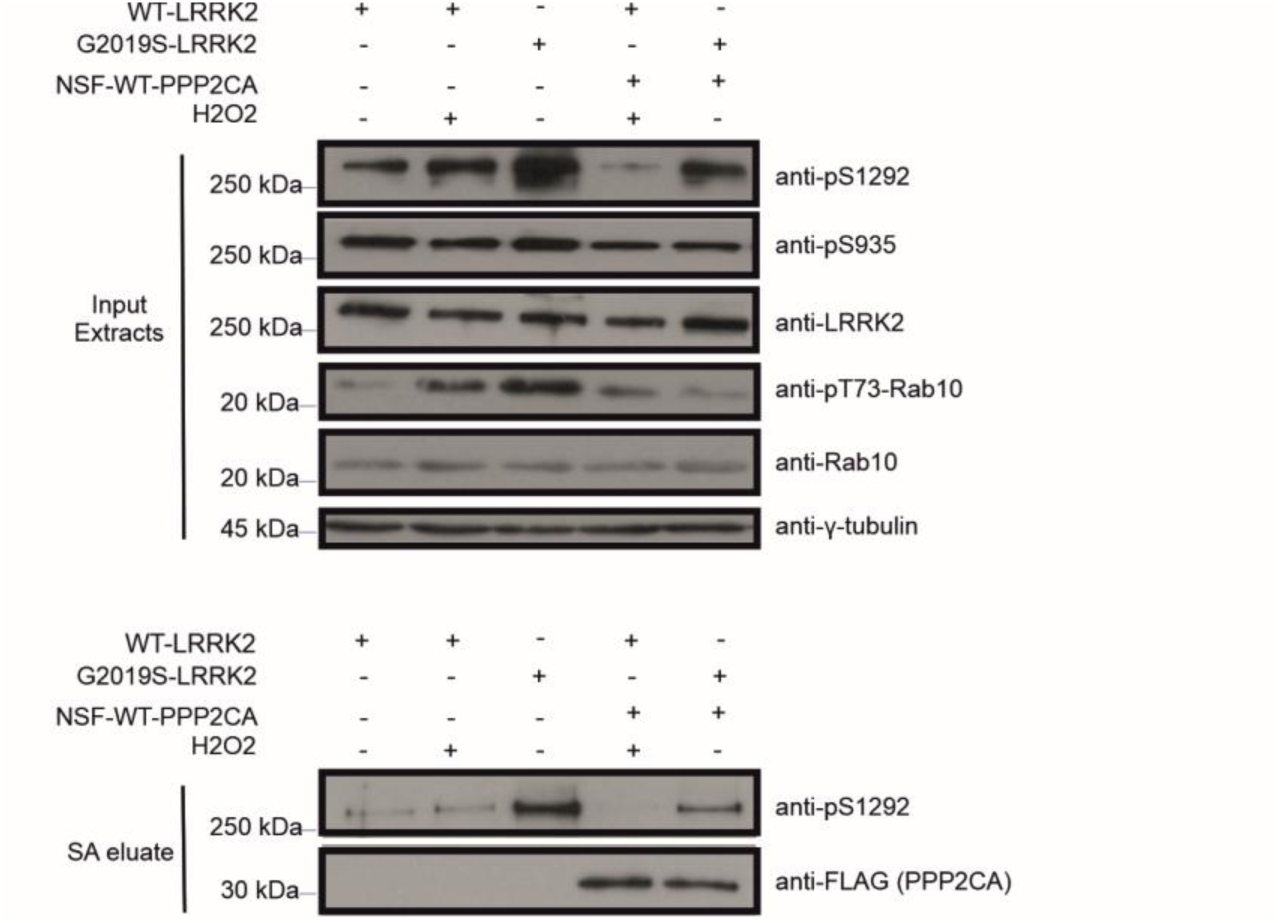
Overexpression of PPP2CA in cells reduces the kinase activity of LRRK2. Co-expression of PPP2CA reduces cellular kinase activity of LRRK2 in HEK293T cells, as assessed by auto-phosphorylation at Ser1292 in both total cell extracts as well as isolated dimers (SA eluate, lower panel), and phosphorylation of Rab10 (T73). Additionally, note that PPP2CA co-elutes with dimeric LRRK2 purified on SA-coated resin.

**Supplementary Fig 2:**
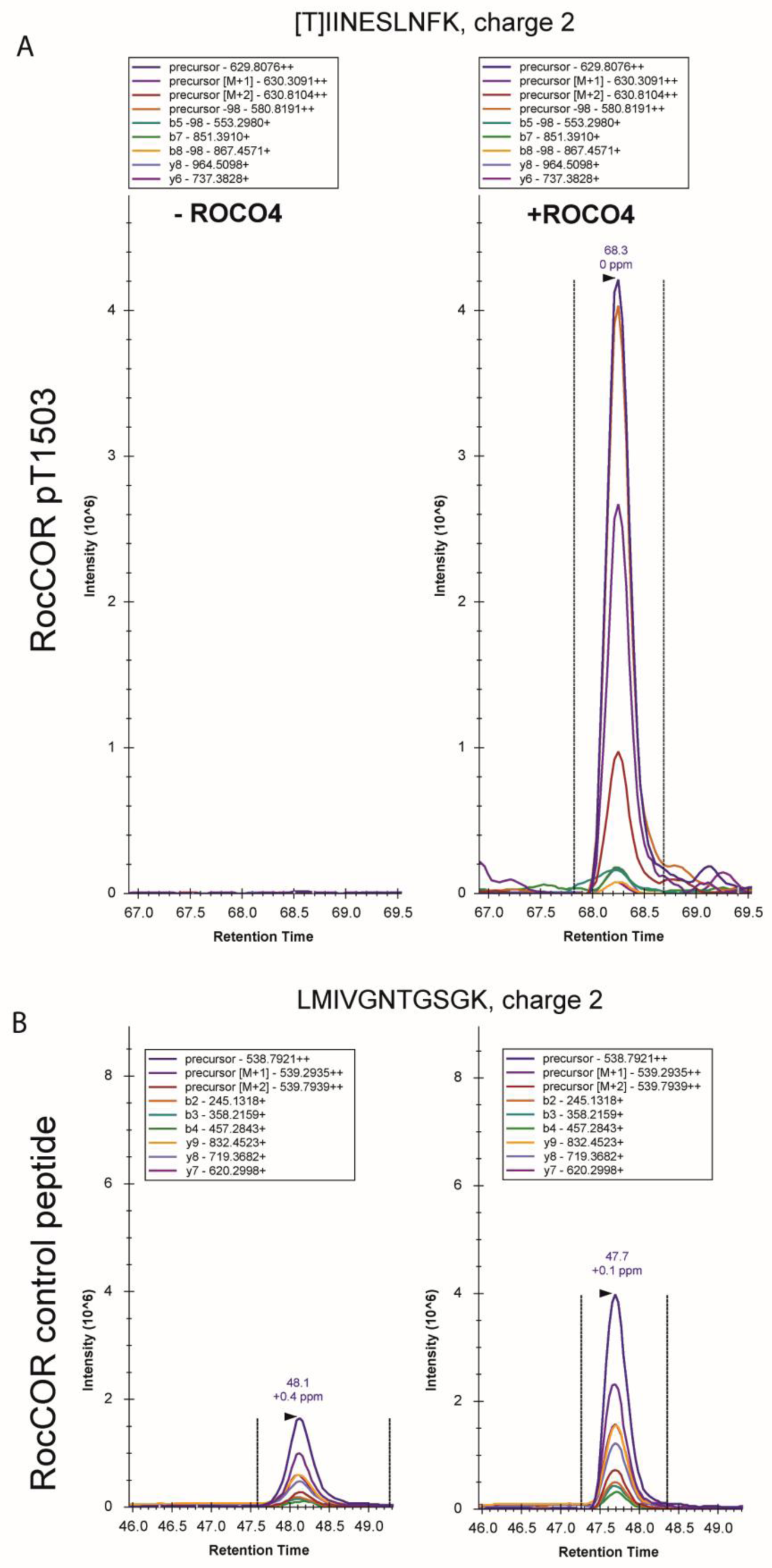
Evidence for LRRK2 pT1503. (A) Extracted PRM spectrum for pT1503 for LRRK2 either incubated without or with ROCO4. (B) Extracted PRM spectrum for a reference peptide corresponding to the LRRK2 Roc sequence. This targeted approach only allowed us to confidently extract the pT1503 from the DIA data.

**Supplementary Fig. 3:**
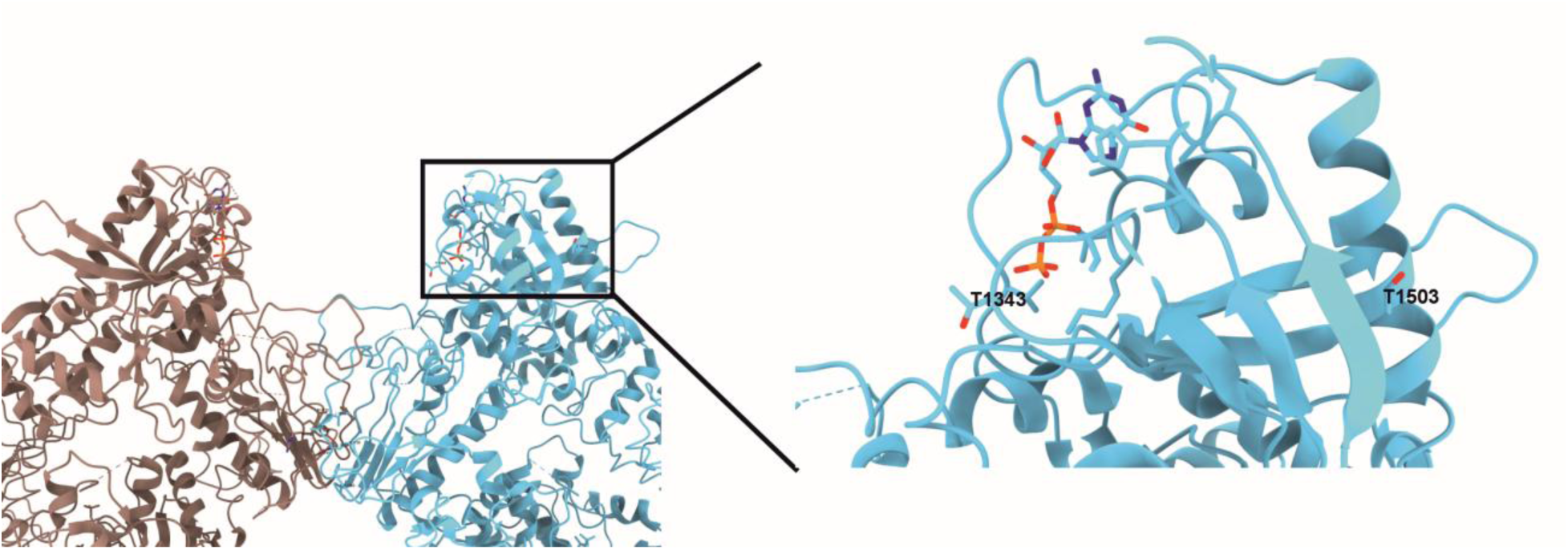
T1503 is in close proximity to the nucleotide-binding pocket of LRRK2. Left: Structure of LRRK2 dimer focusing on the RocCOR dimerization. The two protomers colored brown and blue respectively. Centrally, the dimerization interface can be seen, while on the top right (enlarged) the nucleotide-binding pocket (the T1343 and T1503 residues are found in close proximity to each other and the nucleotide binding pocket. T1343 has been previously shown to affect dimerization and kinase activity of LRRK2 (Gilsbach et al. 2024). PDB code used (7LHT) Myasnikov et al. 2021, Visualised in ChimeraX v. 1.10.

**Supplementary Fig. 4:**
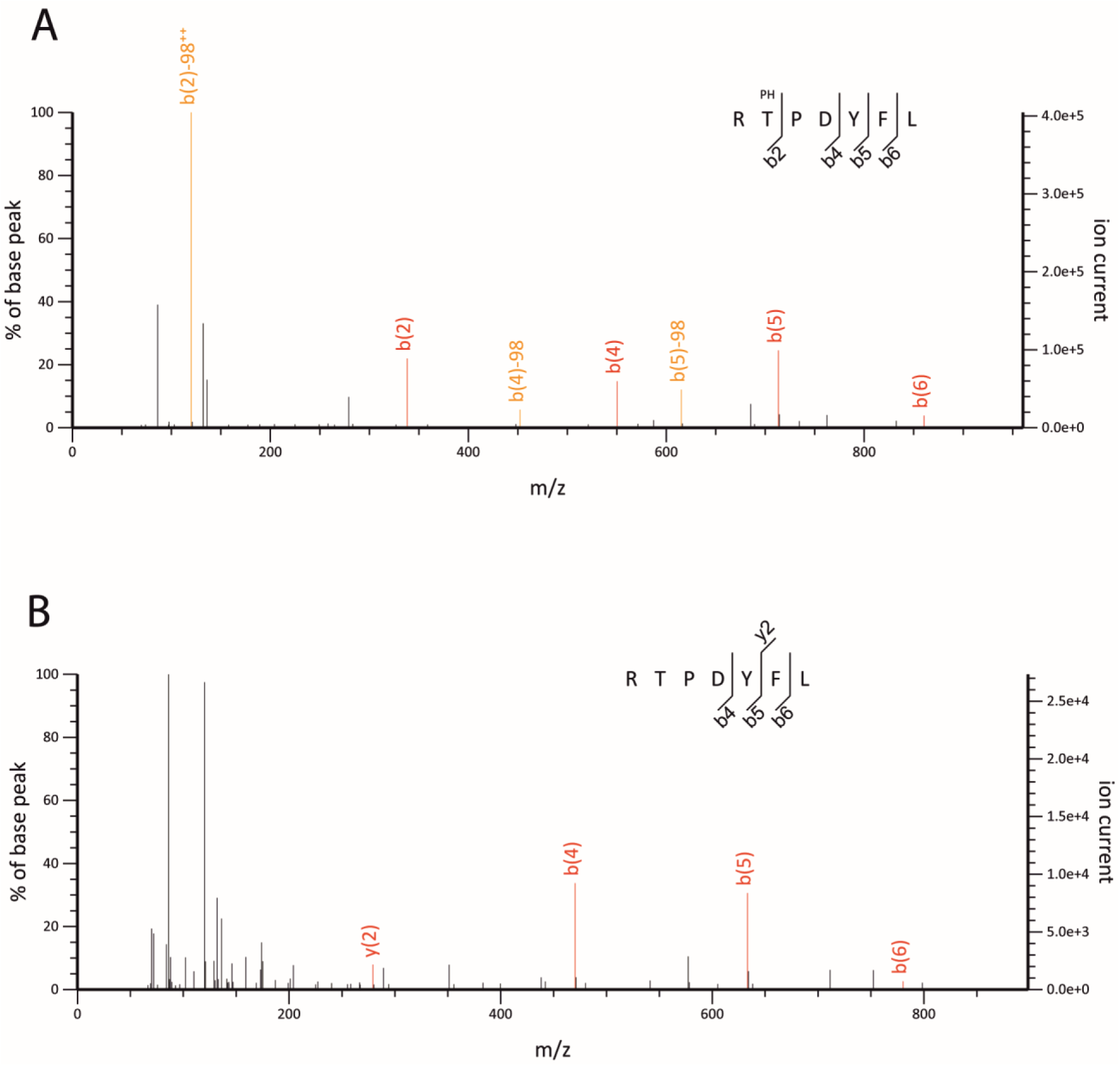
Evidence for the PPP2CA T304 phosphorylation. Mascot-annotated MS2 **s**pectra either for (A) the phosphorylated peptide or (B) unphosphorylated peptide are shown.

**Supplementary Fig. 5:**
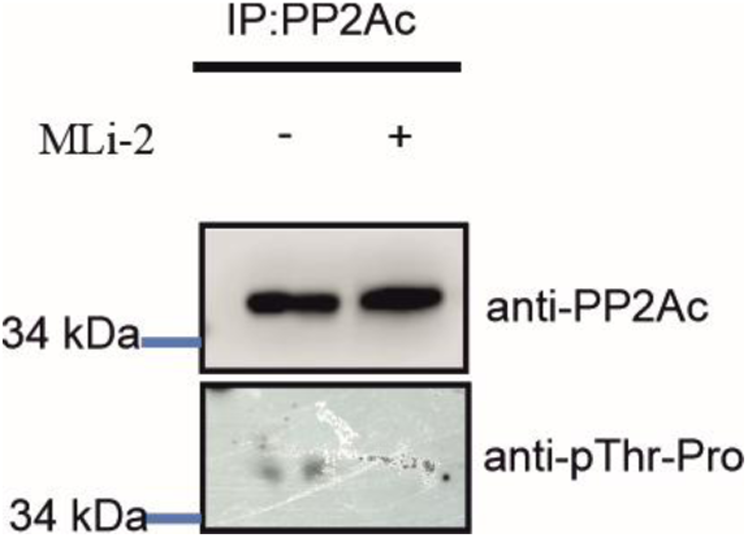
LRRK2 is able to dephosphorylate PPP2CA at T304 residue in A549 cells. A549 lung carcinoma cells were treated with 10nM MLi-2 LRRK2 kinase inhibitor for 90 minutes, prior to cell lysis. Furthermore, the PP2A catalytic subunits were immunoprecipitated. In the absence of the MLi-2 inhibitor, T304 signal of PP2A catalytic subunits is detected, however when the LRRK2 kinase activity is inhibited pharmacologically (by MLI-2), the pT304 signal is diminished.

**Supplementary Fig. 6:**
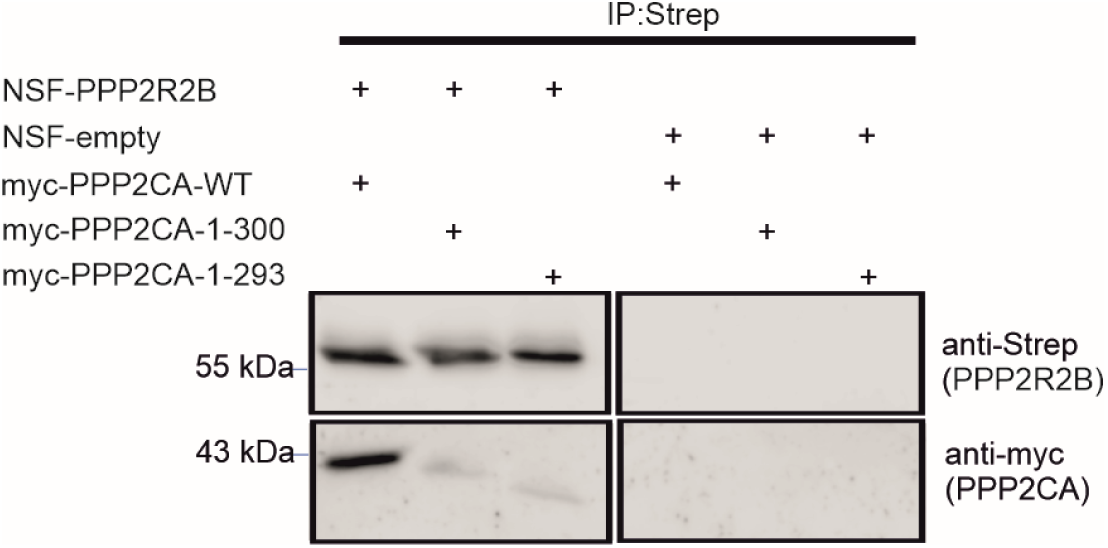
The c-terminal part of PPP2CA is essential for interaction with PPP2R2B subunit. NSF-PPP2R2B or NSF-empty and the indicated myc-tagged PPP2CA constructs were co-transfected in HEK293 cells. Subsequently the cells were lysed and streptavidin beads were used to pull down NSF- PPP2R2B. In addition, we checked whether myc-PPP2CA constructs are in the same complex with NSF- PPP2R2B. While myc-WT-PPP2CA interacts with NSF- PPP2R2B the myc-1-300aa-PPP2CA and myc-1-293aa-PPP2CA proteins are showing a very low interaction affinity with the NSF- PPP2R2B.

